# Genome-wide analysis identifies Homothorax and Extradenticle as regulators of insulin in *Drosophila* Insulin-Producing cells

**DOI:** 10.1101/2022.01.24.477491

**Authors:** M. Winant, K. Buhler, J. Clements, S. De Groef, K. Hens, V Vulsteke, P. Callaerts

**Affiliations:** Laboratory of Behavioral and Developmental Genetics, Department of Human Genetics, KU Leuven - University of Leuven, Herestraat 49, Box 610, B-3000, Leuven, Belgium; Centre for Functional Genomics, Department of Biological and Medical Sciences, Faculty of Health and Life Sciences, Oxford Brookes University, Headington Campus, Oxford, OX3 0BP, UK

## Abstract

*Drosophila* Insulin-Producing Cells (IPCs) are the main production site of the *Drosophila* Insulin-like peptides or Dilps which have key roles in regulating growth, development, reproduction, lifespan and metabolism. To better understand the signalling pathways and transcriptional networks that are active in the IPCs we queried publicly available transcriptome data of over 180 highly inbred fly lines for *dilp* expression and used this as the input for a Genome-wide association study (GWAS). This resulted in the identification of variants in 125 genes that were associated with variation in *dilp* expression. The function of 57 of these genes in the IPCs was tested using an RNAi-based approach. We found that IPC-specific depletion of most genes resulted in differences in expression of one or more of the *dilps*. We then elaborated further on *homothorax*, one of the candidate genes with the strongest effect on *dilp* expression. We found that Homothorax and its binding partner Extradenticle are involved in regulating *dilp2, -3* and *-5* expression and that genetic depletion of both transcription factors leads to phenotypes associated with reduced insulin signalling. Furthermore, we provide evidence that other transcription factors involved in eye development are also functional in the IPCs. In conclusion, we showed that this GWAS approach using gene expression levels as input identified genetic regulators implicated in IPC function and *dilp* expression.

## Introduction

Insulin/IGF signalling (IIS) plays a key role during growth and development but also in regulating reproduction, lifespan, stress resistance and metabolic homeostasis^1–4^. This central role is conserved across metazoans and beyond^5^. Eight Dilps (*Drosophila* Insulin-like peptides) have been described. Dilp1-6 are predicted to bind a single *Drosophila* Insulin Receptor (InR) that results in the activation of the conserved insulin signalling pathway^6^. Dilp7, produced by a group of sexually dimorphic neurons in the ventral nerve cord, acts on the receptor LGR4 and Dilp8, produced in the imaginal discs, binds the receptor LGR3^1,7–10^. Dilp1-3 and 5 are produced in the brain by 14 larval and 16 adult insulin-producing cells (IPCs)^11^. Dilp1 is expressed only after pupation and regulates lifespan and metabolism epistatic to Dilp2^12–14^. The function of Dilp4 is unknown and it is expressed only in the embryonic and larval midgut^1^. Dilp6 is more structurally homologous to mammalian Insulin-like growth factors and is produced in the fat body and glia. It plays an important role in the non-autonomous regulation of the IPCs ^13,15^. Of all Dilps, those most homologous to mammalian insulin and also best studied are Dilp2, -3 and -5, produced in and secreted by the IPCs^16^. In adults, *dilp2* seems to be exclusively expressed in the IPCs, while *dilp3* and *dilp5* are additionally expressed in midgut muscle and in the principal cells of the Malphigian tubules in adults, respectively^17,18^.

The Dilps produced by the IPCs have been shown to be at least partially redundant in their function. Δ*dilp2* mutants, for example, show compensatory increases in *dilp3* and *dilp5* levels. Nonetheless, Δ*dilp2* mutants still have elevated haemolymph trehalose levels not seen in any other single *dilp* mutant, thus showing that the IPC Dilps exhibit independent functions even though they are partially redundant^2,19,20^. Additional evidence for this independent regulation comes from the numerous non-autonomous signals deriving from other tissues that have been shown to regulate the expression or secretion of only a subset of Dilps^21^. The fat body plays a key role in this as it relays the nutritional status of the organism to the IPCs to adjust systemic IIS levels^22^. Low nutritional protein levels, for example, induce the release of Eiger from the fat body into the haemolymph. Eiger binds to and activates its receptor Grindelwald on the IPCs to inhibit *dilp2* and *dilp5* expression but not *dilp3*^23^. Neuronal input differentially affects *dilp* expression. Knockdown of the octopamine receptor OAMB yielded increased *dilp3* expression while leaving *dilp2* and *dilp5* levels unaltered, while knockdown of the 5-HT1a serotonin receptor results in increased *dilp2* and *dilp5* and unaltered *dilp3*^24^. The autonomous factors acting in the IPCs that mediate this independent regulation remain poorly understood.

To identify novel autonomous regulators of IPC *dilp* expression we use the *Drosophila* Genome Reference Panel (DGRP) consisting of more than 200 highly inbred fly lines whose genomes have been sequenced. The DGRP has been used effectively to identify genes that determine many quantitative phenotypes such as starvation resistance^25^, aggressive behavior^26^, mushroom body morphology^27^, cocaine consumption^28^ and lifespan^29^. While previous studies relied on quantitative behavioural or physiological parameters, we used the variation in *dilp2, -3* and *-5* mRNA levels of 183 (DGRP)^25,30^ fly lines to identify genetic variants that correlate with *dilp2, -3* and *-5* expression levels. Genes associated with these variants were validated for a role in determining *dilp2, -3* and *-5* levels in the IPCs with an RNAi-based approach. We then elaborated on the function of one of the strongest candidate genes, the homeobox transcription factor Homothorax (Hth). We show that it is an important regulator of *dilp2, -3* and *–5* expression in the IPCs and is necessary for normal systemic insulin signalling. We then show that the Hth interactor Extradenticle (Exd) is equally required in the IPCs. Based on the previously established roles of Ey, Dac and now Hth/Exd in regulating both IPC and eye field development we then hypothesized that other members of the Retinal Determination Gene Network (RDGN) could also be active in the IPCs^31–33^. We show that most RDGN genes are expressed in the IPCs and regulate *dilp2 and -5* expression. Our results further unravel the transcriptional regulatory networks that control *dilp* expression and implicate a reshuffled RDGN being active in the IPC to control insulin expression.

## Results

### GWAS based on expression levels for *dilp2,3* and *5* reveals sexually dimorphic putative regulators

We collected whole body expression data for *dilp2, -3 and -5* of 183 DGRP lines^30^ (Figure 1 A; Supplemental File 1). Considerable interline variation can be noted and significantly higher read counts were detected in males compared to females for the three *dilps*. We then ran a GWA analysis for each of the *dilps* and identified a total of 271 variants that were associated with at least one *dilp* and in one sex (Figure 1B). These 271 variants correspond to 125 unique genes (Figure 1 B). Based on RNA-seq data of the adult IPCs (Supplemental File 2) to ensure that genes were expressed in the adult stage and RNAi line availability we assessed the role of 57 candidate genes (Supplemental file 3) in regulating *dilp2 -3* and -5 expression in the IPCs using qPCR. This showed that 50 out of 57 had significant effects on *dilp2, -3* or *-5* regulation in one or both sexes (Figure 1 C).

**Figure 1:**
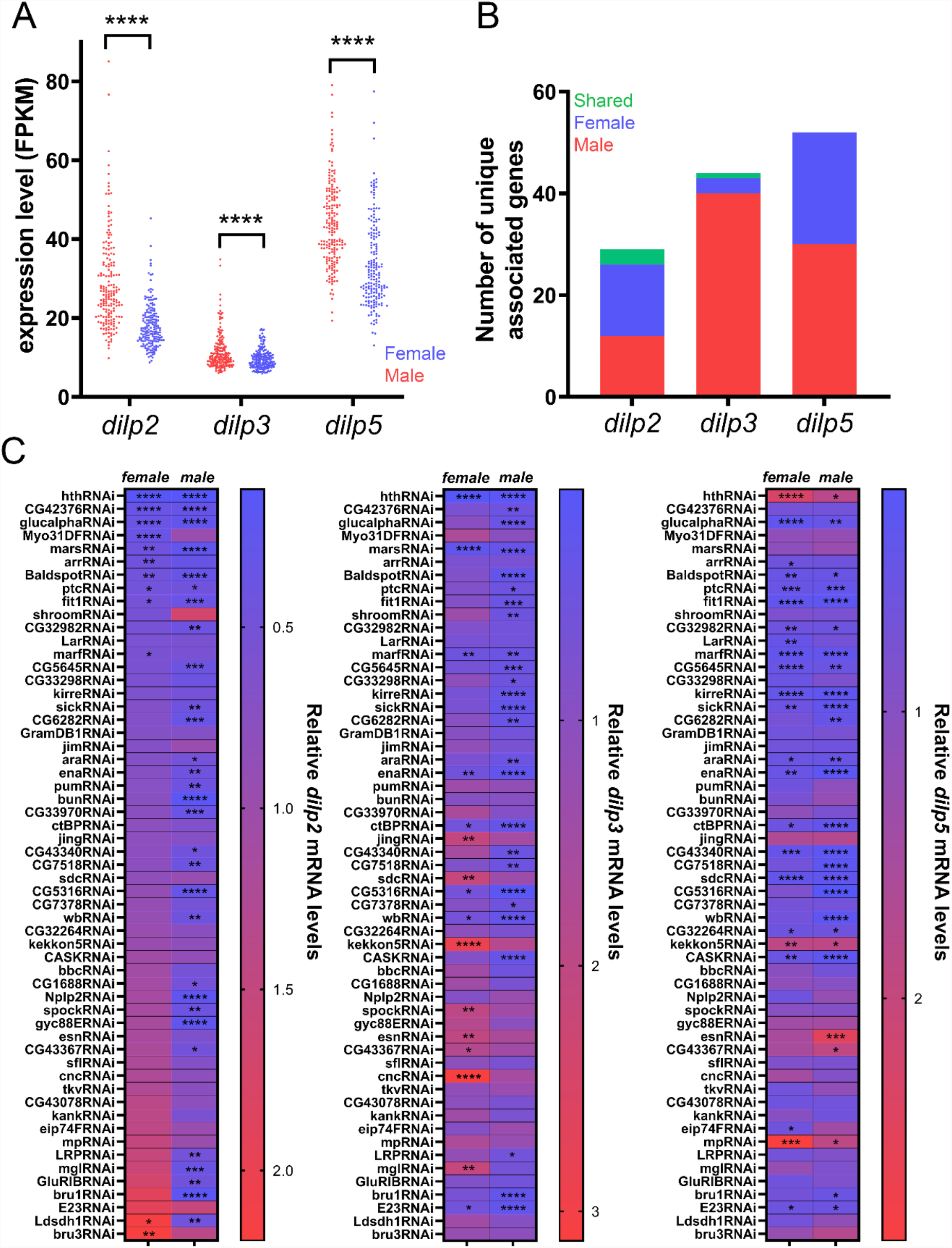
DGRP Screen and validation. (A) Mean *dilp2, -3 and dilp5* FPKM levels were collected from Huang et al.^30^ and are significantly higher for males than females for *dilp2* (p < 0.0001, unpaired Welch’s T-test, n=183), *dilp3* (p < 0.0001, unpaired Welch’s T-test, n = 182) and *dilp5* (p < 0.0001, student’s T-test n = 182). (B) Number of unique genes associated for each dilp/sex (male dilp2: 12, female dilp2: 14, shared dilp2: 3, male dilp3: 40, female dilp3: 3, shared dilp3: 2, male dilp5: 30, female dilp5: 22 and shared dilp5: 0). (C) Mean qPCR levels for *dilp2, -3 and dilp5* upon RNAi-mediated knockdown using *Dilp2-GAL4*^*R*^*;btlGAL80* (values are means of the relative mRNA quantity of n=3 consisting of 10-15 fly heads each). Results of female *dilp2* levels were used to sort the RNAi lines (low to high). (* p < 0.05, ** p <0.01, *** p<0.001, **** p < 0.0001, Dunnett’s multiple comparisons compared to an *mCherry*^*RNAi*^ control line, which itself had no effect on *dilp* expression (Figure S2))

Most genes that were identified in the GWAS were identified in only one sex suggesting a highly sexually dimorphic regulation of *dilp* expression (Figure 1 B). However, when comparing changes in expression of the *dilp*s upon knockdown of the 57 candidate genes in the IPCs we noticed that the *dilp* levels in both sexes showed significant correlation (Figure 2 A-C). To better understand this apparent contradiction with the GWAS results, we classified per *dilp* each candidate gene as sexually dimorphic if RNAi mediated depletion in the IPCs resulted in a significant reduction or increase of *dilp* mRNA levels in one sex only, or if it had an opposite effect on *dilp* levels in both sexes. The latter possibility was only seen for *Ldsdh1* where knockdown resulted in increased *dilp2* levels in females and decreased *dilp2* levels in males (Figure 2 A’). Overall, we found that out of 57 candidate genes, knockdown of 34 genes resulted in changes in *dilp2* expression of which 79% (27) were sexually dimorphic (Figure 2A’). Changes in expression of *dilp3* were seen for 34 out of 57 candidate genes with 76% sexually dimorphic (Figure 2B’). Finally, 30 out of 57 genes resulted in altered expression of *dilp5* with 30% being sexually dimorphic (Figure 2C’). Of note, the genes that result in changes in *dilp* expression upon knockdown are not the same for each *dilp*, consistent with the hypothesis that there is significant independent regulation of transcription for the *dilps*. In conclusion, we identify significant sexually dimorphic effects for regulation of *dilp2* and *dilp3* and less for *dilp5*.

**Figure 2:**
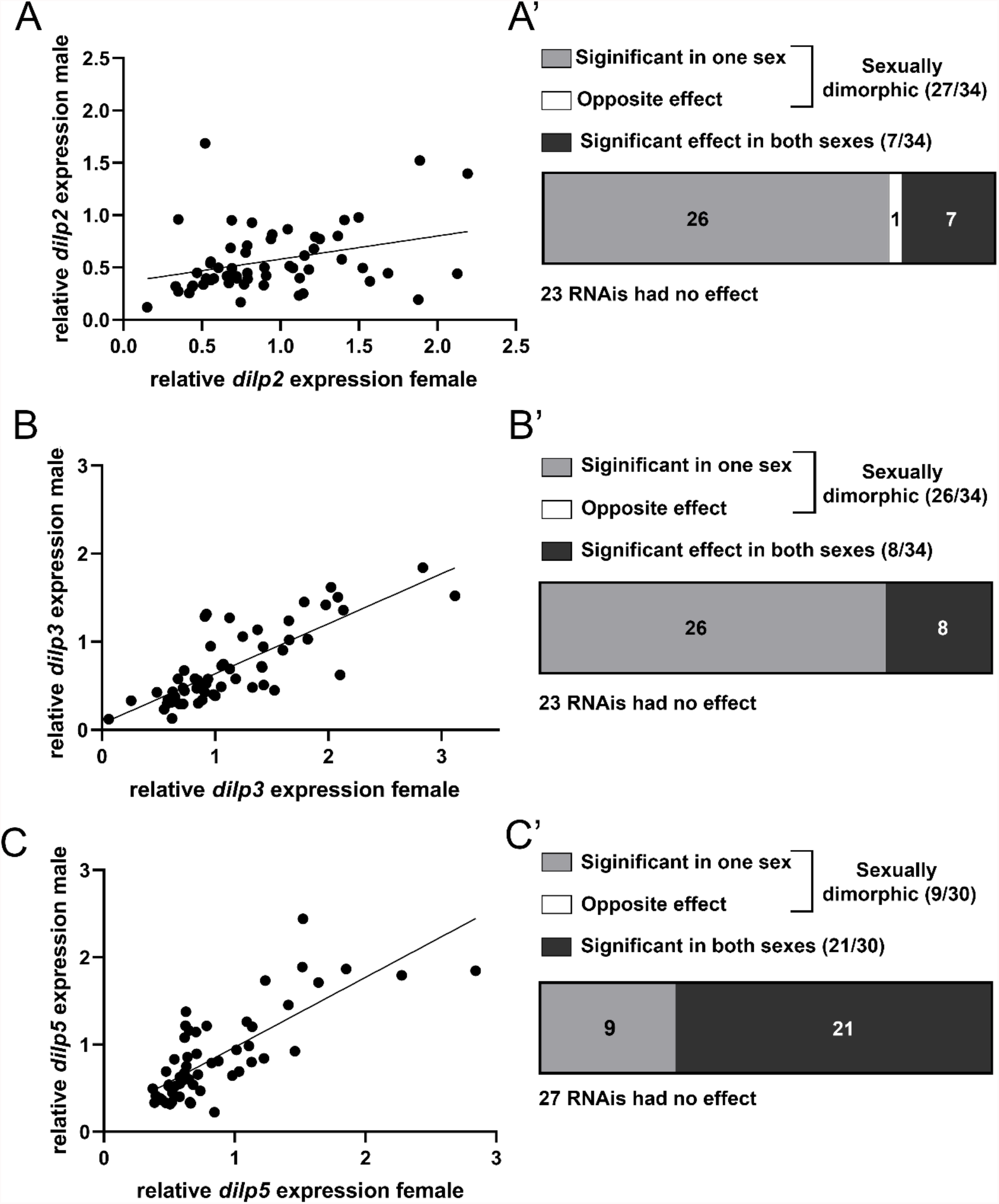
Knockdown of candidate genes in the IPCs produces correlated effects on *dilp* expression between the sexes. (A) The mean relative quantity of *dilp2* upon RNAi-mediated knockdown in the IPCs between female and male flies is mildly correlated (Spearman r = 0.3467, p = 0.0081). (A’) Out of 57 tested genes, 27 had sexually dimorphic effects on *dilp2* expression, one (*Ldsdh1*) having opposite effects on *dilp2* expression. 30 genes had no effect (23) or the same effect (7) on *dilp2* levels. (B) Relative dilp3 between the sexes are highly correlated. (Spearman r = 0.7955, p < 0.0001) (B’) 26 genes had a sexually dimorphic effect on *dilp3* expression, 31 had no effect (23) or the same effect (8). Relative quantity levels of *dilp5* are correlated (Spearman r= 0.7699, p < 0.0001). (C’) 9 genes had a sexually dimorphic effect on *dilp5* levels while 48 genes had no effect (27) or the same effect (21) on *dilp5* expression.

### Homothorax knockdown reduces *dilp2* and *-3* mRNA expression and Dilp2,3 and 5 protein levels

The strongest decline in both *dilp2* and *dilp3* levels in males and females was noted for the transcription factor Homothorax. IPC-specific downregulation resulted in significant downregulation of both *dilp2* and *dilp3* mRNA levels while *dilp5* mRNA levels were significantly increased (Figure 1C). First, we validated the expression of Hth in the IPCs of L3 larvae and of adults by using antibodies directed against Hth (Figure 3 A – B). We then elaborated on the effect of Hth on *dilp* expression by characterising the effect on Dilp protein levels by using specific antibodies against Dilp2, -3 and -5^34^. When compared to sibling controls this revealed a strong reduction of Dilp2 -3 and -5 immunoreactivity within the IPCs (Figure 3 C-E). Similar results were found upon *hth* knockdown in the larval IPCs (Figure S3 A-C). Knockdown of Homothorax, however, did not have a significant effect on adult body weight (Figure S2 D).

**Figure 3:**
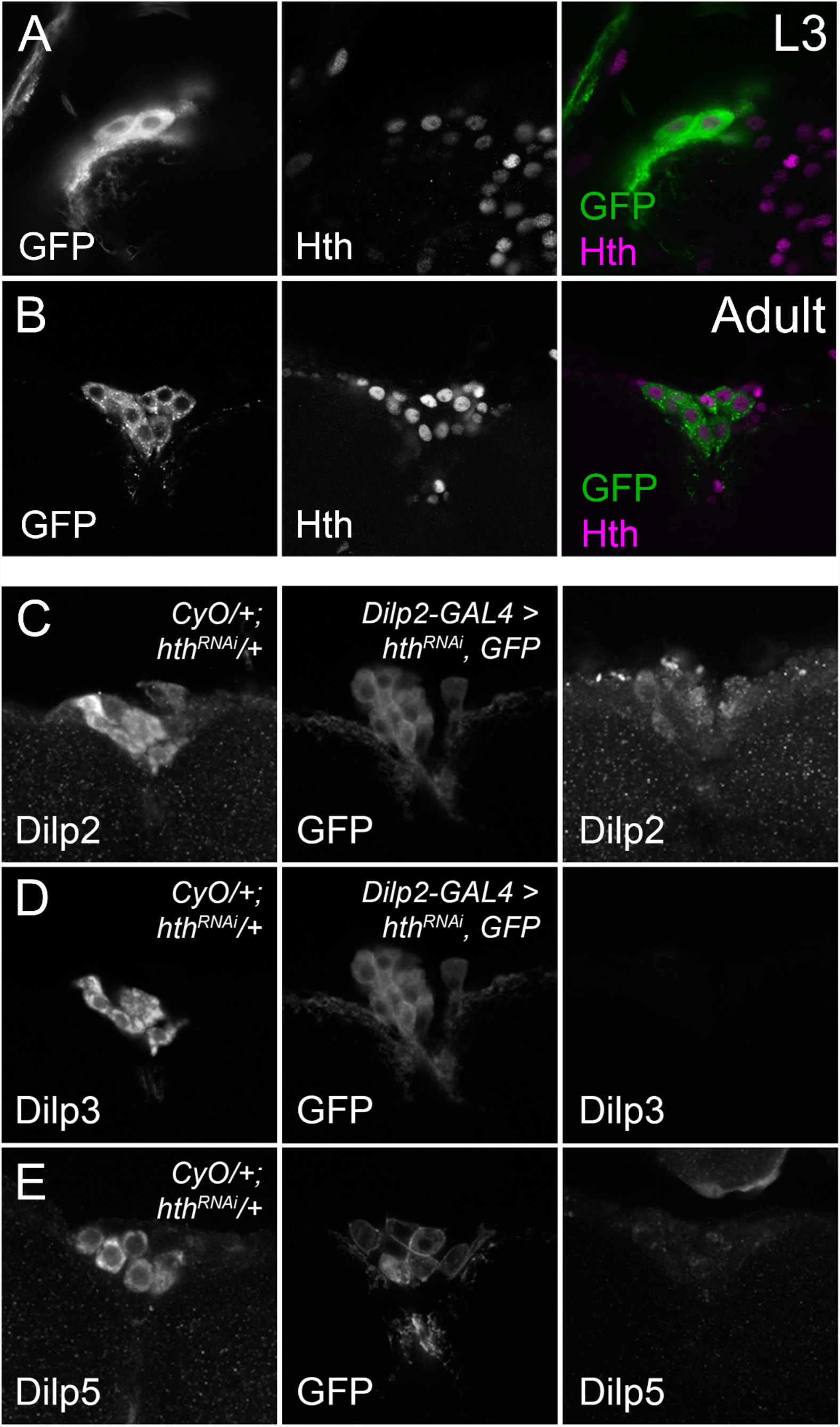
Hth is expressed in larval and adult IPCs and is necessary for normal Dilp protein levels. Antibody staining detects Hth in L3 (A) and adult (B) nuclei in IPCs that are marked by CD8-GFP expression. Knockdown of *hth* reduces Dilp2 (C), Dilp3 (D) and Dilp5 (E) protein expression in the IPCs compared to balanced age-controlled siblings reared in identical environmental conditions. Brains from knockdown and balanced siblings were dissected and stained in the same tube. Expression of CD8-GFP was used to distinguish control and test genotypes, and to label the IPCs. Images were acquired with identical confocal settings and representative examples are shown.

### Knockdown of the Hth cofactor, Extradenticle, phenocopies Hth depletion

During development, Hth physically interacts with the homeobox transcription factor Exd. Exd forms a heterodimer with Hth, and is necessary for nuclear localization of Hth^35^. An α-Exd antibody detects protein expression in both L3 and adult IPCs (figure 4 A - B). We therefore hypothesized that Exd, together with its co-factor Hth, regulates *dilp* expression in the IPCs. Knockdown of *exd* strongly reduced expression of all *dilps* with two different RNAi lines, virtually identical to the effects of *hth* knockdown (Figure 4 C-E). Dilp protein levels were also reduced in both L3 larval and adult stages, although Dilp5 protein could be detected in wandering L3 stage (Figure S3 A - F). We then checked the effects of Hth and Exd depletion on oxidative stress resistance, which is regulated by systemic insulin signalling^36^. Genetic depletion of Hth and Exd in the IPCs strongly increased oxidative stress resistance consistent with a reduction of systemic insulin signaling. This effect was even stronger than for Ey, another transcription factor that positively regulates *dilp* expression (Figure 4 F).

**Figure 4:**
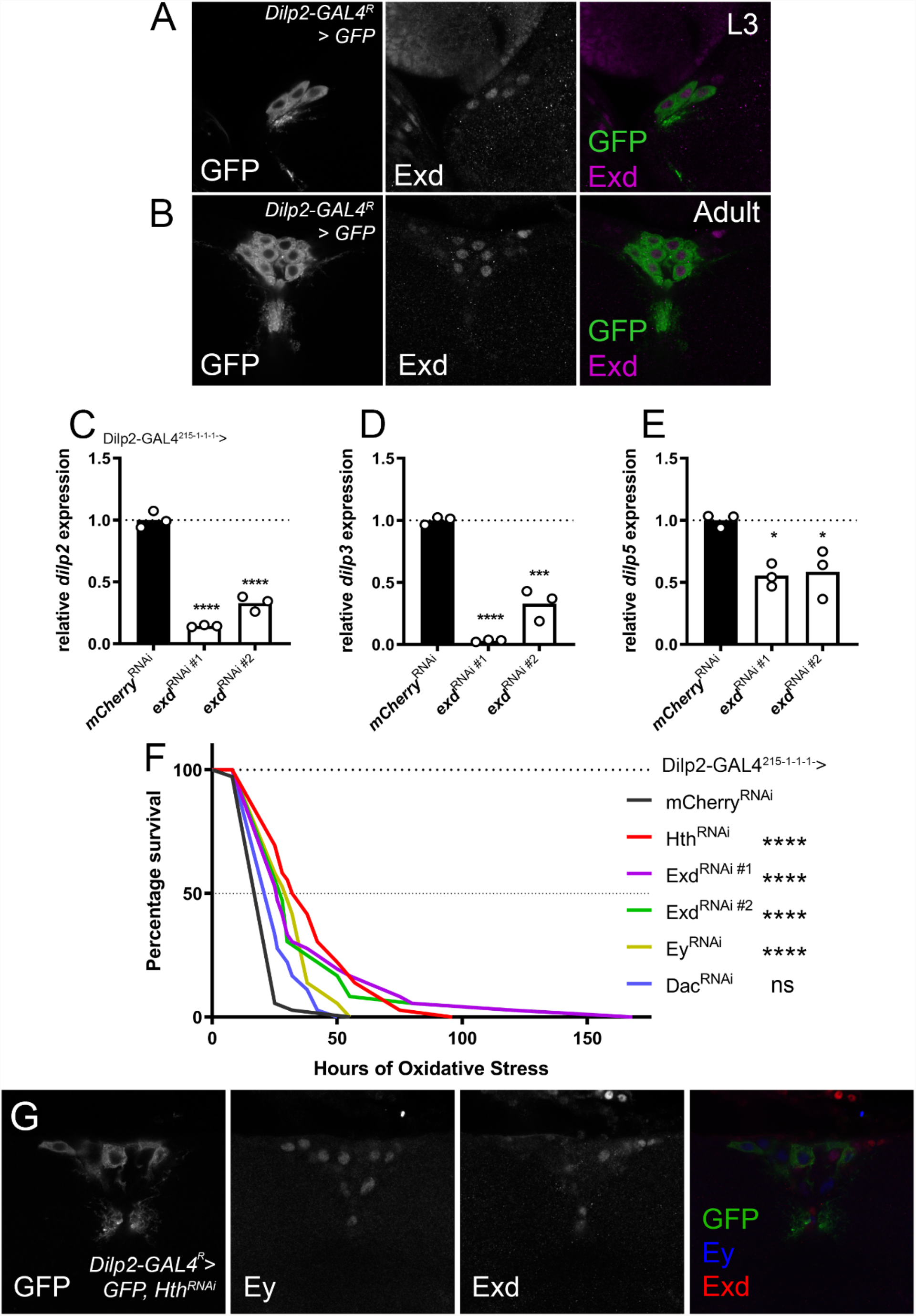
The Hth interactor Exd, is expressed in the IPCs and is equally required for *dilp* expression. (A-B) Antibodies against Exd detect expression in L3 and adult IPC nuclei, marked by *dilp2-GAL4*^*R*^ driven *UAS-CD8GFP expression* (C-E) Knockdown of *exd* using two different RNAi lines in the IPCs using *dilp2-GAL4*^*215-1-1-1*^ reduced expression of *dilp2, dilp3* and *dilp5*. Flies were reared at controlled densities. RNA was collected from 10 female fly heads of flies aged between 12-15 days old. After normalization, *dilp* expression was relativized to a control *dilp2-GAL4*^*215-1-1-1*^ driving *UAS-mCherry*^RNAi^. (F) Hth, Exd and Ey depetion increases lifespan in oxidative stress conditions. Flies were reared on normal food and aged to 3 days old and placed on food containing 20mM paraquat at controlled densities. Mortality was scored regularly (*hth*^*RNAi*^, *exd*^*RNAi*^ #1, *exd*^*RNAi*^ *#2, ey*^*RNAi*^ p <0.0001, *dac*^*RNAi*^ not significant, log-rank test with Bonferroni correction for multiple testing). (G) In IPCs genetically depleted of *hth*, expression of Ey is unaffected, though Exd is not detectable in IPC nuclei.

### Genes of the Retinal Determination Gene Network regulate *dilp* expression in the IPCs

In the mammalian pancreas, the Hth/Exd homologs Meis1/PBX1/2 function upstream of PAX6/Dach1/2 where they are required to induce *pax6* expression^37,38^. We thus hypothesized that Hth would have a similar regulatory relationship with Ey in the IPCs. Depletion of *hth* in the IPCs did not result in differences in Ey levels in the IPCs, while Exd expression was no longer detectable in the IPC nuclei, showing that Hth is required for normal Exd localization and/or expression, but not for normal Ey expression (Figure 4 G).

A well-established gene regulatory network in which Hth/Exd act is the retinal determination gene network (RDGN), necessary for eye development^33^. We and others had already described a role for other RDGN members: Eyeless (Ey) and Dachshund (Dac) in regulating the IPCs. Therefore, we hypothesized that a version of this regulatory network controls *dilp* expression in the IPCs^31,32^ To test this, we first queried the transcriptome of both larval and adult IPCs (Cao et al^39^, and this study) for expression of 11 different RDGN members. Of these 11 TF, all were expressed to some degree in the adult IPCs, with *hth, gro, optix, dac, eya, ey* and *dan* showing expression in both stages. (Figure 5A). These results were validated with antibody and transgenic reporter lines where available (Figure 5B - Q). Only Eyg and Eya showed no detectable expression in the IPCs but the overall staining quality for Eyg and Eya was poor (Figure 5 N-Q) and while RNA-seq results for the larval stage suggested that only a subset of RDGN TF were expressed in L3 IPCs, all except Eyg and Eya are clearly detectable at the protein level using immunohistochemical reagents. This shows that most transcription factors of the RDGN are expressed in the IPCs. To determine whether these transcription factors are active in regulating *dilp* expression in the IPCs, we genetically depleted these TFs using *Dilp2-GAL4*^*215-1-1-*1^ and their respective RNAi lines^40^. Except for *dac* and *toe*, knockdown of these TFs resulted in a significant reduction in *dilp2* and *dilp5* expression compared to D*ilp2-GAL4*^*215-1-1-1*^ flies driving a control RNAi under identical environmental conditions (Figure 5 R - T). Knockdown of *dac* reduced *dilp5*, and increased *dilp2* levels, while knockdown of *toe* only reduced *dilp5* expression. Furthermore, only the depletion of *tsh* and *ey* affected *dilp3* levels (Figure 5 S). This suggests that the RDGN is re-used in the regulation of IIS in the IPCs.

**Figure 5:**
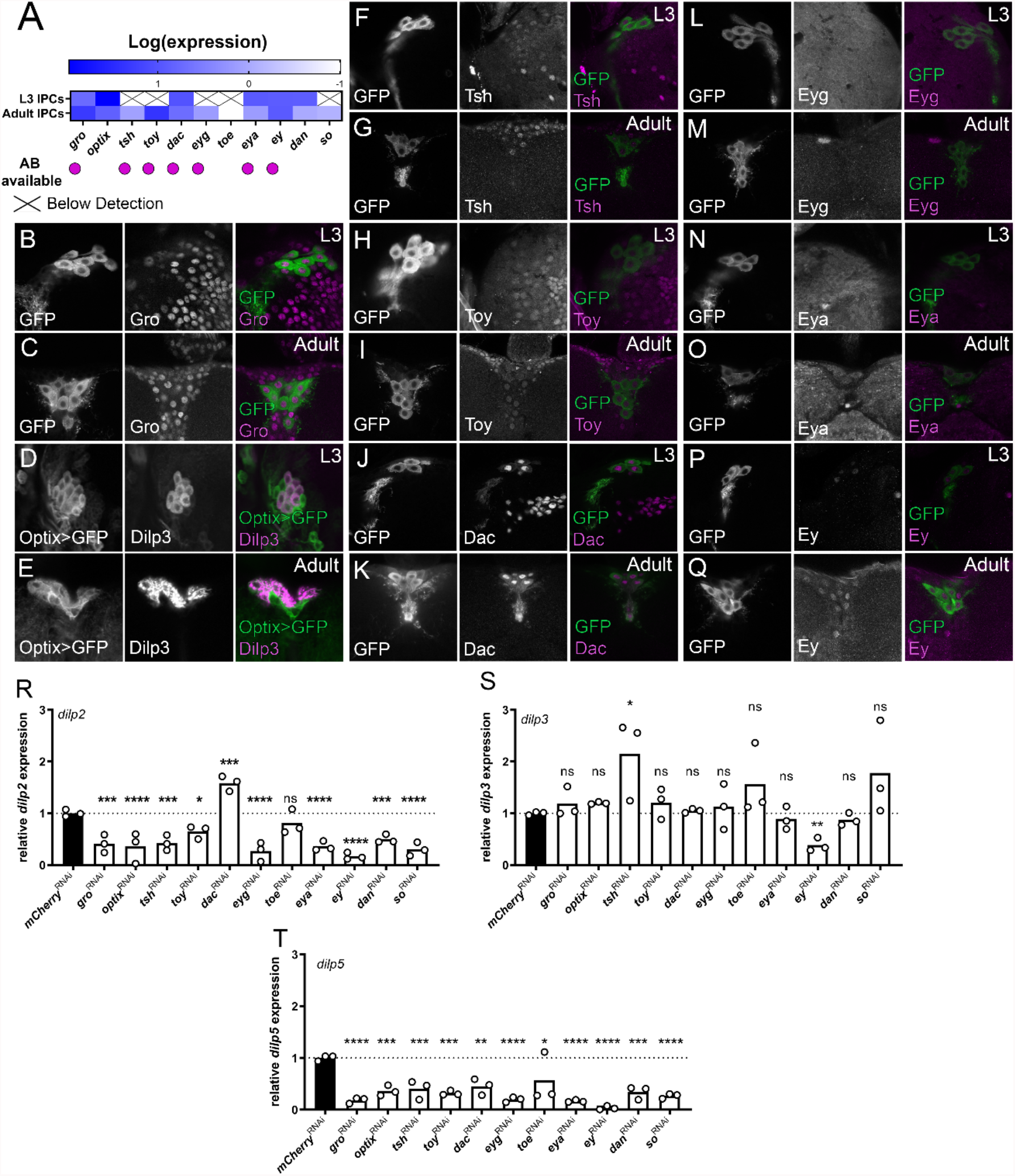
Most RDGN transcription factors are expressed in the IPCs and are required for normal adult *dilp* expression. (A). Additional RDGN transcription factors are expressed in larval and/or adult IPCs (Larval transcriptome data is taken from Cao et al., (2014)^39^ while adult transcriptome data was generated for this study. (B-Q) Gro, Tsh, Toy, Dac and Ey are readily detected in L3 and adult IPCs, only Eyg and Eya exhibited no detectable protein expression in the IPCs. Adult *Dilp2-GAL4*^*R*^ *> CD8GFP* flies were reared to sexual maturity and brains were dissected and stained with available ABs directed against RDGN TFs. Expression of CD8GFP was used to label IPCs. For Optix, an Optix-GAL4 driver drove expression of the CD8GFP and was localized with an antibody against Dilp3, which labelled the IPCs. (R-T) Genetic knockdown of all RDGN TFs in the IPCs using *Dilp2-GAL4*^*215-1-1-1*^ except *dac* and *toe* resulted in significant reduction of *dilp2* expression (one-way ANOVA with post-hoc Dunnett’s test comparing to *mCherry*^*RNAi*^ controls, n = 3), *tsh* depletion in the IPCs increases dilp3 expression (one-way ANOVA with post-hoc Dunnett’s test comparing to *mCherry*^*RNAi*^ controls, n = 3) and all were required for dilp5 expression (one-way ANOVA with post-hoc Dunnett’s test comparing to *mCherry*^*RNAi*^ controls, n = 3). Flies were reared at controlled densities and RNA was collected from 10 female fly heads of flies controlled for age and diet.

In summary, these data show that a repurposed RDGN is used in the IPCs. Knockdown of *hth* and *exd* affects all three Dilps. On the other hand, most RDGN TFs regulate either *dilp2* and *dilp5*, while *toe* regulates only *dilp5* and *tsh* regulates all three. Our data provide an interesting example of how regulatory networks are re-used in other developmental contexts to achieve different transcriptional outcomes.

## Discussion

The IPCs are key regulators of growth and metabolism in the fly but the regulatory networks governing their development and function remain poorly understood. In this study, we use the variation in *dilp* expression across DGRP lines to identify genes that are necessary for normal autonomous *dilp* regulation. There was very little overlap between the sexes for the identified candidate genes (3 genes for *dilp2*, 2 genes for *dilp3* and none for *dilp5*). Furthermore, only one gene was shared between the *dilps* (*Glucalpha* for *dilp3* and *dilp5* in females). The results from our RNAi based approach to validate the role of these putative regulators revealed that 50 out 57 tested impacted expression of at least one of the *dilps* and that in many cases sexually dimorphic effects were noted (Figure 1 C, Figure 2 A-C’). Additionally, genes implicated in the regulation of one *dilp* often had additional effects on the other dilps. Variants in Hth, for example, were identified to correlate with *dilp5* levels in males but not females. Depletion of Hth in the IPCs produced a pronounced effect on *dilp5* levels and additionally resulted in a strong reduction of both *dilp2* and *dilp3* levels in both sexes (Figure 1 C). Several possible explanations for this discrepancy exist. One explanation is that our RNAi-based approach might have obfuscated the effect of the specific variants which might have sex-specific and/or *dilp*-specific effects. The compensatory effects previously noted in *dilp* regulation might add an additional layer of complexity impacting our observations^2,41^. Furthermore, the identified genes might also have additional roles in other tissues and thus also contribute to regulating dilp expression in a non-autonomous fashion^21^.

Our approach revealed that Hth is required in the adult IPCs to regulate *dilp* expression (Figure 1 C). IPC depletion of Hth resulted in almost complete absence of *dilp2* and *dilp3* mRNA in both sexes (~10% of control) while doubling *dilp5* mRNA levels. Immunohistochemistry confirmed the strong reduction of Dilp2 and Dilp3 at the protein level but an increase on Dilp5 protein levels was not seen. The latter could be due to increased Dilp5 secretion or possibly also a failure to produce properly matured Dilp5 protein (Figure 3 C-E). Further analysis revealed that Hth regulates *dilp* expression with its binding partner Exd. Exd depletion had effects on *dilp* expression and increased oxidative stress resistance comparable to that observed upon Hth depletion. The effects were stronger than those for the transcription factor Ey which also serves as an important regulator of IPC development and *dilp* expression^42^ (Figure 4 C-F). In the mammalian pancreas the fly homolog of Ey, Pax6, is required to establish the cellular fate of endocrine cell types and to promote expression of their respective hormone gene products. To do this, Pax6 cooperates with Dach1/2, a relationship also observed in the Ey/Dac regulation of *dilp5* in *Drosophila*^32^. Upstream of Pax6, the Meis gene family members Meis1/2/Prep1/2 and Pbx1/2 are required to promote pancreatic *pax6* expression^37^. Depletion of Hth in the IPCs, however, leaves Ey expression unaffected while resulting in the absence of Exd in IPC nuclei (Figure 4 G). This shows that while the TFs are identical between flies and humans, their regulatory relationships are not. Similarly, differences in regulatory relationships are also seen during eye development where Meis1/2 regulate *pax6* in the mouse lens. In *Drosophila*, however, Hth has no direct regulatory relationship with *ey* or its paralog *toe*^43,44^. Key TFs are thus conserved in *Drosophila/*mammalian eye development and IPC/pancreas development while the regulatory relationships between the TFs have diverged during evolution.

We further show that the genes of the RDGN are active in the IPCs to regulate insulin signalling, mostly via control of *dilp2* and *dilp5* expression. Of all tested TFs only Ey and Hth/Exd are necessary for promoting *dilp3* expression while Tsh depletion resulted in an upregulation of *dilp3* levels (Figure 5 S). Other RDGN TFs were not required to regulate *dilp3*, and most regulated *dilp2* and *dilp5*. In literature, co-regulation between *dilp2* and *-5* is commonly reported, despite them having distinct functions^19^. However, given the complex regulatory relationships between the RDGN TFs in the developing eye, it is surprising to see that, in general, in the IPCs their regulatory relationship is largely positive, promoting expression of these two *dilps* (Figure 6 A). One explanation is that since a couple of the RDGN TFs are already known to be expressed at the neuroblast stage^45^, depleting these TFs during development might result in a failure to properly establish IPC cell fate. Additionally, a more complex regulatory architecture may exist that is relevant in other physiological contexts to which the *dilps* are sensitive, such as in changing dietary conditions, aging, or oxidative stress^46^. Of note are the opposing effects of Dac and Ey on *dilp2* expression, while still physically interacting to promote *dilp5*. This implies the presence of *dilp*-specific regulatory relationships. We do, however, not provide evidence for the direct regulation of the *dilps*, but we hypothesize, based on the strong effects of Hth/Exd on *dilp* expression that they may directly regulate the *dilps*, particularly *dilp2* and *dilp3*, while others may contribute via indirect interaction or regulation of additional factors (Figure 6B).

**Figure 6:**
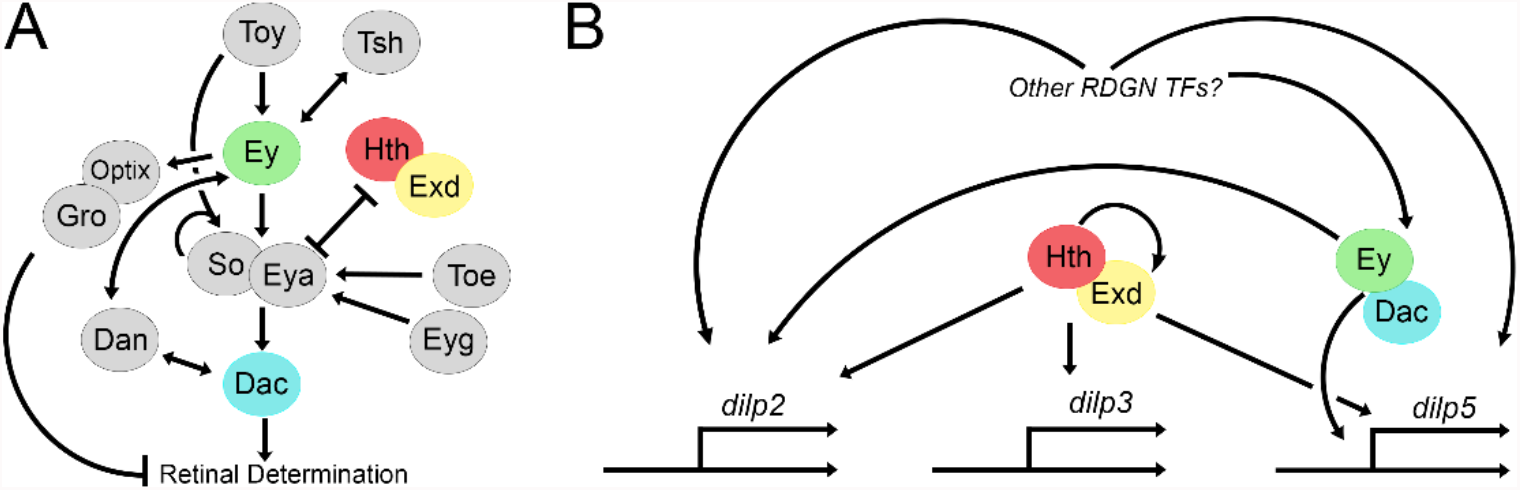
A reshuffled RDGN regulates *dilp* expression in the IPCs. (A) Schematic showing the regulatory relationships of the RDGN in the developing eye (modified from Kumar^33^ and Bürgy-Roukala *et al*.^55^). (B) Speculative model of the RDGN in *dilp* regulation. Hth and Exd as a pair directly or indirectly regulate *dilp2, -3 and -5* expression, while Ey and Dac directly regulate *dilp5* expression^32^ and via other factors *dilp2* and *dilp3*. Other RDGN TFs directly or indirectly regulate *dilp2* and *dilp5 expression*.

Interestingly, a study by Okamoto *et al*.^32^, which originally identified Dac as a co-regulator of *dilp5* together with Ey, also knocked down a number of other RDGN TFs (*optix, eya, eyg, tio, so, toy*). They find small and often not significant effects on *dilp* levels when compared to our results. We speculate that differences between our results and theirs lay in the age of the flies, the different *dilp2-*GAL4 line that was used or the differences in dietary conditions. Another study based on type 2 diabetes risk loci in humans by Peiris *et al*.^47^, showed that IPC-specific knockdown of *Optix* resulted in decreased circulating Dilp2 levels. Our results suggest that this reduction in Dilp2 release might be indirect due to the reduction of *dilp2* mRNA levels upon *Optix* depletion.

To conclude, our approach identified many novel regulators of *dilp* expression with a prominent role for Hth/Exd in regulating *dilp* expression in both sexes. In addition, we show that the RDGN is repurposed to regulate the development and function of the IPCs. The RDGN is thus a paradigm of how evolution uses the finite genetic tools available to establish the structural and functional diversity observed in biology.

## Methods

### Fly Stocks and Transgenes used in this Study

Drosophila stocks were reared at 25°C on Nutrifly (Genesee Scientific) for the initial DGRP validation screen. Subsequent experiments were performed using standard yeast fly food recipe (See Table 1, below). Stocks were ordered from the Bloomington Drosophila Stock Center (BDSC) or Vienna Drosophila RNAi Center (VDRC). The following GAL4 lines were used: *Dilp2-GAL4R; btl-GAL80* (*BDSC# 37516*, described in^16^) and Dilp2-GAL4^215-1-1-1 40^.To visualize IPCs, *Dilp2-GAL4* lines controlled expression of *UAS-CD8GFP* (BDSC# 5137). A list of the RNAi lines used in the DGRP screen are included in Supplemental File 3. RNAi lines for the RDGN follow-up experiments include *hth-RNAi (BDSC# 34637), ey-RNAi (BDSC# 32486), dac-RNAi(BDSC# 35022), exd-RNAi #1 (BDSC# 34897), exd-RNAi #2 (BDSC# 29338), tsh-RNAi (BDSC# 35030), optix-RNAi (BDSC# 55306), toy-RNAi (BDSC# 29346), toe-RNAi (BDSC# 50660), dan-RNAi (BDSC# 51501), eyg-RNAi (BDSC# 26226), gro-RNAi (BDSC# 35759), so-RNAi (BDSC# 35028)* and *eya-RNAi (BDSC# 35725)*.

### Flyfood recipe

| Ingredient | Amount |
| --- | --- |
| Water | 15.2 L |
| Cornmeal | 1000 g |
| Yeast | 180 g |
| Agar | 112.5 g |
| Dextrose | 270 g |
| Molasses | 600 mL |
| Propionic Acid | 150 mL |
| 10%Hydroxybenzoate | 22.5 g |
| 99.8% Ethanol | 225 mL |

### Candidate gene identification

GWA analyses on *dilp* values were performed using the DGRP2 webtool (http://dgrp2.gnets.ncsu.edu/) ^25,48^. Mean FPKM values of *dilp2, -3* and *-5* were used as the input and genes associated with variants that reached a nominal significance threshold of p < 10^-3 in the mixed effects model in one of the sexes were considered for further validation. Input files are provided in Supplemental File 1.

### Transcriptome data

IPCs were fluorescently labelled with dilp2-GAL4; UAS-mCD8GFP. GFP-positive cells were manually sorted according to Nagoshi et al^49^. In short, 100 brains were dissected from adult flies, 3-5 days after eclosion into ice-cold dissecting solution (9.9 mM HEPES-KOH buffer, 137 mM NaCl, 5.4 mM KCl, 0.17 mM NaH2PO4, 0.22 mM KH2PO4, 3.3 mM glucose, 43.8 mM sucrose, pH 7.4) containing 50 µM d(−)-2-amino-5-phosphonovaleric acid (AP5), 20 µM 6,7-dinitroquinoxaline-2,3-dione (DNQX), 0.1 µM tetrodotoxin (TTX), and immediately transferred into modified SMactive medium (5 mM Bis-Tris, 50 µM AP5, 20 µM DNQX, 0.1 µM TTX) on ice. Brains were digested with l-cysteine-activated papain (50 units ml−1 in dissecting saline) for 20 min at 25 °C. Digestion was quenched with a fivefold volume of the medium, and brains were washed twice with the chilled medium. Brains were triturated with a flame-rounded 1,000-µl pipette tip with filter followed by a flame-rounded 200-µl pipette tip. Single, GFP-positive cells were selected, then reselected a further 2 times until approximately 100 fluorescent cells were pooled. Total RNA was extracted using a Nucleospin RNA micro kit (Macherey-Nagel). cDNA synthesis and adapter ligation were performed with the SMARTer cDNA synthesis kit (Takara Bio). 51bp paired-end reads were generated on an Illumina HiSeq2000 sequencer. Resulting reads were submitted to the European Nucleotide Archive, Accession Number PRJEB50165.

Quality assessment and read trimming was performed with fastp (v. 0.23.1)^50^ with default settings. Reads were then pseudo-aligned to the reference Drosophila genome (BDGP6.32) and TPM values determined using salmon (v. 1.5.0)^51^ and summarized to the gene level using tximport (v. 1.18.0)^52^

### RNA extraction and qPCR

For the validation of candidates from the DGRP, offspring was collected after eclosion and maintained at controlled densities of 20 total flies, aged to 3-5 days and snap frozen on dry ice. 10 to 15 fly heads were used per replicate for RNA extraction. For subsequent qPCR experiments concerning the RDGN transcription factors, flies were aged until 12-15 days old and only female flies were used for qPCR experiments.

Total RNA was isolated using TRI™ reagent (Invitrogen) and reverse transcription was performed on 1µg RNA using Transcriptor First Strand Synthesis kit (Roche). Primer sequences used are listed in Table S1. qPCR was performed on a Step-One-Plus using the SYBR Green detection system. Transcript levels were normalized to Rp49 transcript levels (initial screen) or using the geometric means of RpS13 & Rp49 (transcription factor experiments). Mean ΔCt values were statistically compared as described below. Relative quantitation of transcript levels to control genotypes were calculated using the ΔΔCt method and plotted with GraphPad Prism software to visualize expression differences.

### Primers used in this study

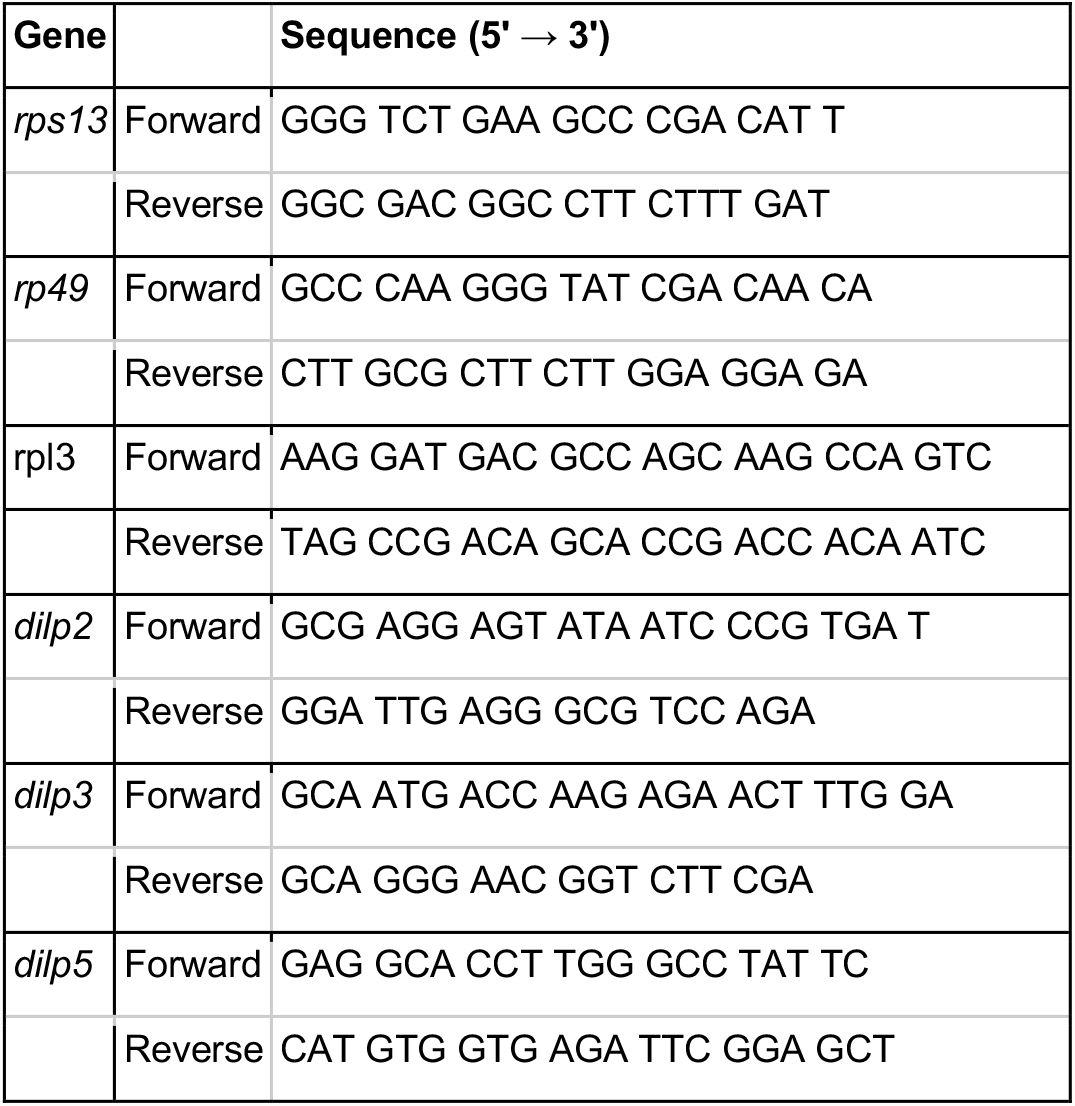

### Immunohistochemistry

Adult and larval brains were dissected in 1x PBS prior to 30 minutes fixation with 4% formaldehyde. Primary antibodies were incubated overnight at 4°C in PAXD (PBS containing 1% BSA, 0.3% Triton X-100, 0.3% deoxycholate). Antibodies used in this study were mouse α-GFP (University of Iowa, Developmental Studies Hybridoma Bank (DSHB) 8H11; 1:100), rabbit α-GFP (Life Technologies, Ref#: A6455; 1:1000), rabbit α-Hth (Dr. Richard Mann, 1:200), mouse α-Ey (DSHB E4H8, 1:5), rabbit α-Exd (Dr. Richard Mann, 1:200), mouse α-Dac (DSHB mAbdac2-3, 1:20), α-Gro (DSHB anti-Gro, 1:10), rat α-Tsh (Dr. John Bell, 1:25), rat α-Toy (1:50), guinea pig α-Eyg (Dr. Natalia Azpiazu, 1:100), mouse α-Eya (DSHB eya10H6, 1:100), rat α-Dilp2 (Dr. Pierre Leopold, 1:1000), rabbit α-Dilp3 and rabbit α-Dilp5.

Rabbit antibodies directed against the Dilp3 partial peptide sequence DEVLRYCAAKPRT and against the Dilp5 peptide sequence RRDFRGVVDSCCRKS were generated as a service by Thermo Fischer Scientific Inc. For immunostaining, α-Dilp3 antibodies were used at a dilution of 1:100 and α-Dilp5 antibodies used at a dilution of 1:400. Secondary antibodies used include goat α-mouse FITC & Cy3 and goat α-rabbit FITC & Cy3 (Jackson Immunoresearch). All secondary antibodies were used at a dilution of 1:200. Samples were mounted in Vectashield® Mounting Medium (Vector Laboratories Cat #H-1000). Immunohistochemistry images were taken with an Olympus FluoView FV1000 confocal microscope and processed using ImageJ64 (1.6.0_65; FIJI) ^53,54^ and Photoshop software.

### Oxidative stress resistance with Paraquat

Female flies were collected and aged until 12-15 days old, then transferred onto fly food containing 20mM Paraquat (Sigma 75365-73-0). Deaths were scored regularly and plotted in GraphPad Prism. Median lifespan was compared using a log-rank test with post-hoc Bonferonni correction for multiple testing.

### Determination of Adult Fly Weight

Adult flies 3-7 days after eclosion, experimental (*Dilp2-GAL4;btlGAL80 > UAS-hth-RNAi, UAS-CD8GFP*) and control siblings (*CyO; btlGAL80 > UAS-hth-RNAi, UAS-CD8GFP*) reared in identical environmental conditions were anesthetized using chloroform and weighed on a Mettler Toledo XS204 scale (Wet Weight; d = 0.1 mg). Mean weight between experimental and control siblings was compared using an unpaired Student’s t-test with GraphPad software, and the relative change in adult weight between experimental and control genotypes were plotted.

### Female fecundity

Eclosed females were collected and allowed to mate for 48 hours with sibling males in a 4 female to 1 male sex ratio. Females were then collected and placed in groups of 4-5 in separate vials. Flies were transferred onto fresh food daily and the number of eggs was counted for 14 days.

### Statistics

All statistics were conducted using the GraphPad Prism software. All statistics were performed on raw data after normalization, where applicable (i.e. qPCR). For assays comparing means, unpaired T-tests for pairwise comparisons or one-way analysis of variance (ANOVA) tests for multiple comparisons were used. To test whether variance significantly differed between samples, an F-test was performed for pairwise comparisons and both Brown-Forsythe and Bartlett’s Tests performed for multiple comparisons. If the standard deviations between samples were significantly different, a T-test with Welch’s correction (Welch’s T-test) was performed for multiple comparisons, whereas the Geisser-Greenhouse correction was always applied on all one-way ANOVA analyses, as sphericity of the data was not assumed. When comparing means to a single control sample, Dunnett’s multiple comparisons test was used. For lifespan analysis under oxidative stress, median lifespan was compared with a log-rank test, with Bonferroni correction for multiple testing. For all statistical analyses, we assumed a significance level of 0.05.

## Figure Legends

**Supplementary Figure 1:**
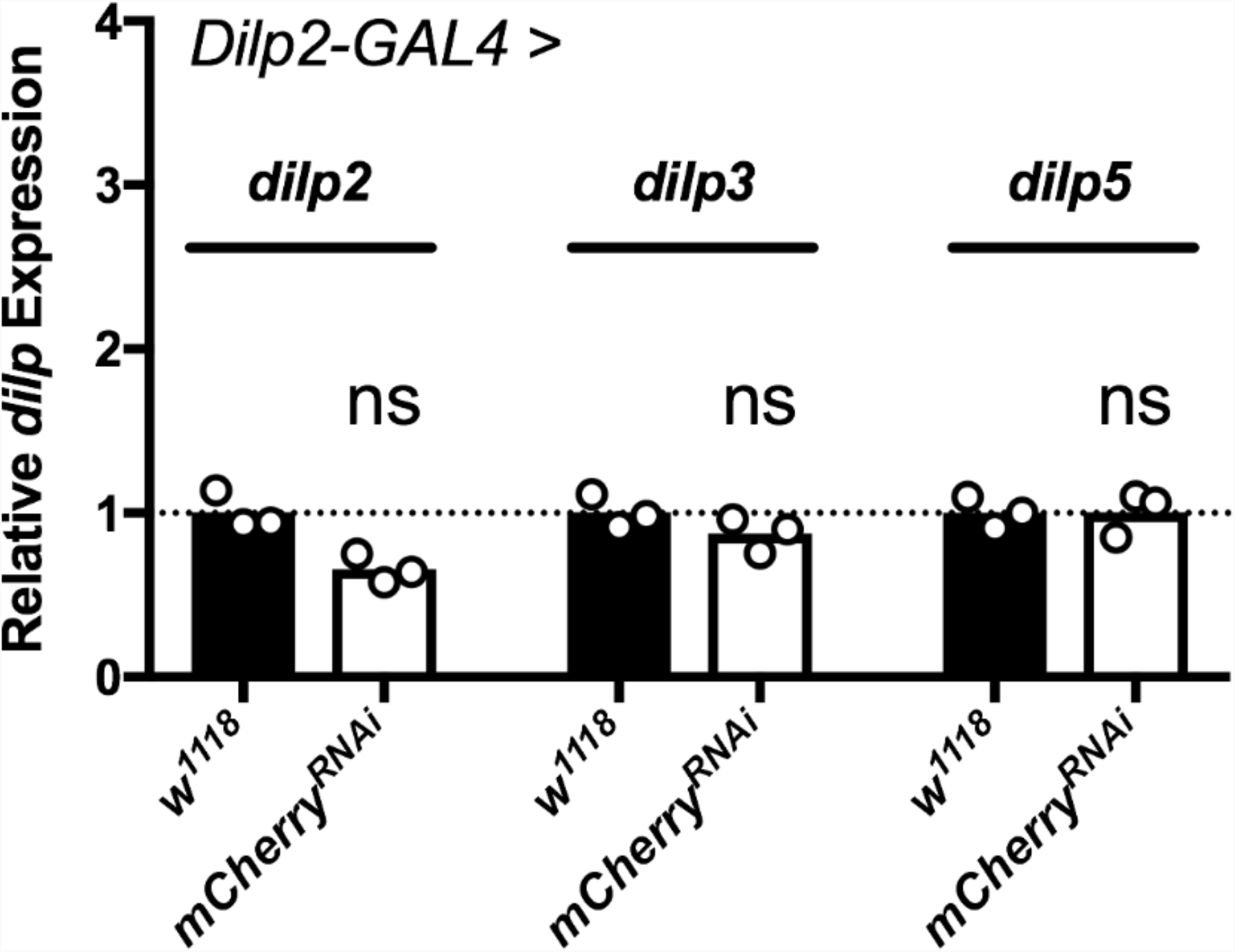
Expressing an *mCherry*^*RNAi*^ in the IPCs does not affect *dilp* expression. *dilp2, -3* and *-5* mRNA levels are not significantly different in *Dilp2-GAL4*^*R*^ > *mCherry*^*RNAi*^ compared to *Dilp2-GAL4*^*R*^ */+* fly heads (unpaired two tailed Student’s t-test p > 0.05).

**Supplementary Figure 2:**
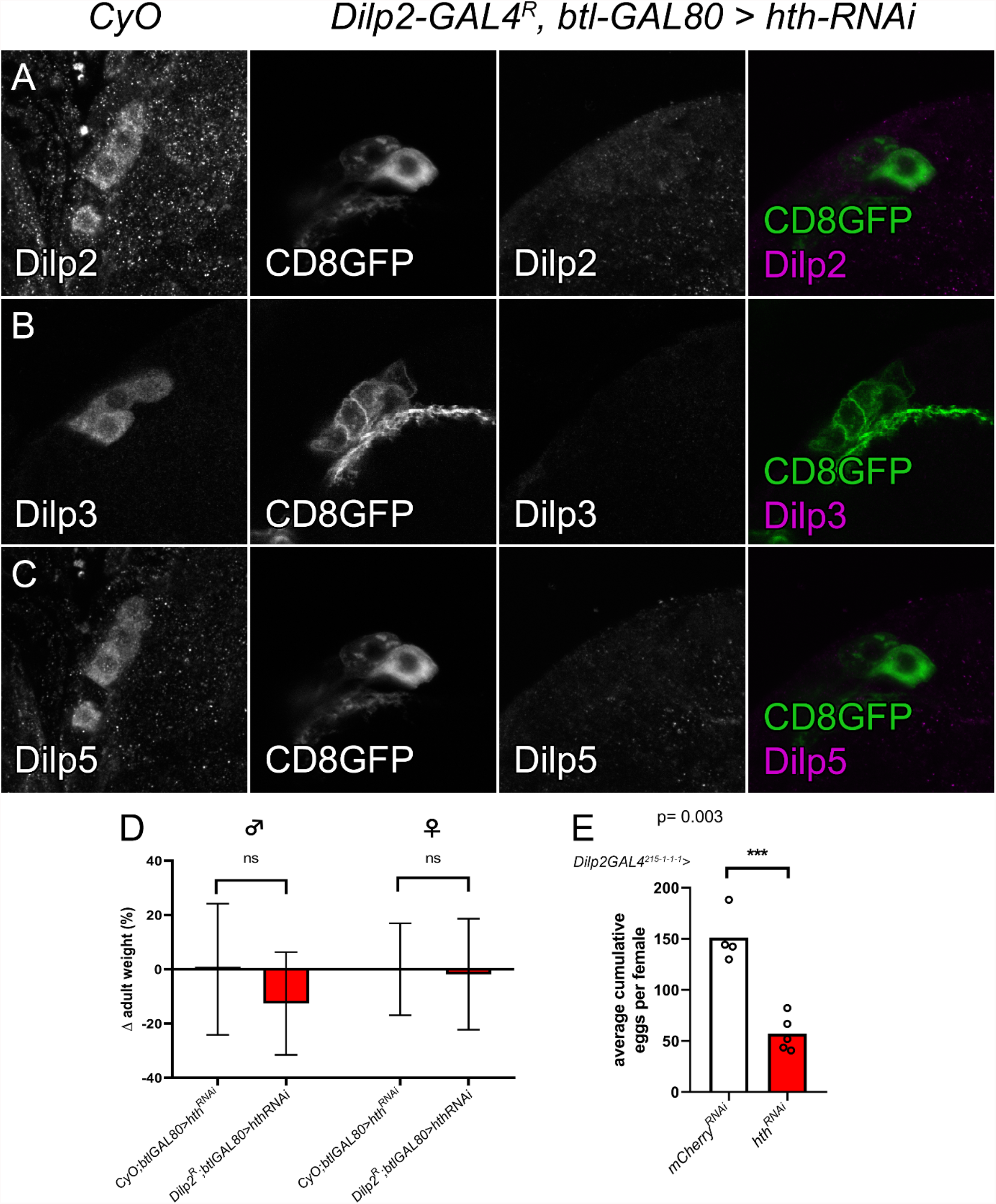
Knockdown of *hth* in the larval IPCs reduces Dilp2, -3 and -5 protein levels with no clear effect on systemic growth. (A-C) Genetic depletion of Hth in the IPCs using *Dilp2-GAL4*^*R*^*;btl-GAL80* resulted in reduced (A) Dilp2, (B) Dilp3 and (C) Dilp5 protein levels compared to control siblings reared in identical conditions. Flies were reared and age-controlled as described above, then brains from knockdown and balanced sibling flies were dissected and stained in the same tube. Expression of *CD8GFP* was used to differentiate control and test genotypes, and label IPCs. Images were acquired using identical confocal settings. (D) No significant differences in adult fly weight upon *hth* knockdown in the IPCs. Adult fly weight was compared to control siblings reared in identical nutritional and crowding conditions, mean weights were compared with unpaired two tailed Student’s t-test (p > 0.05). (E) IPC depletion of *hth* resulted in decreased female fecundity, every data point is the average (4-5 females per vial) cumulative number of eggs layed for 14 days (p = 0.003, unpaired two tailed Student’s t-test).

**Supplementary Figure 3:**
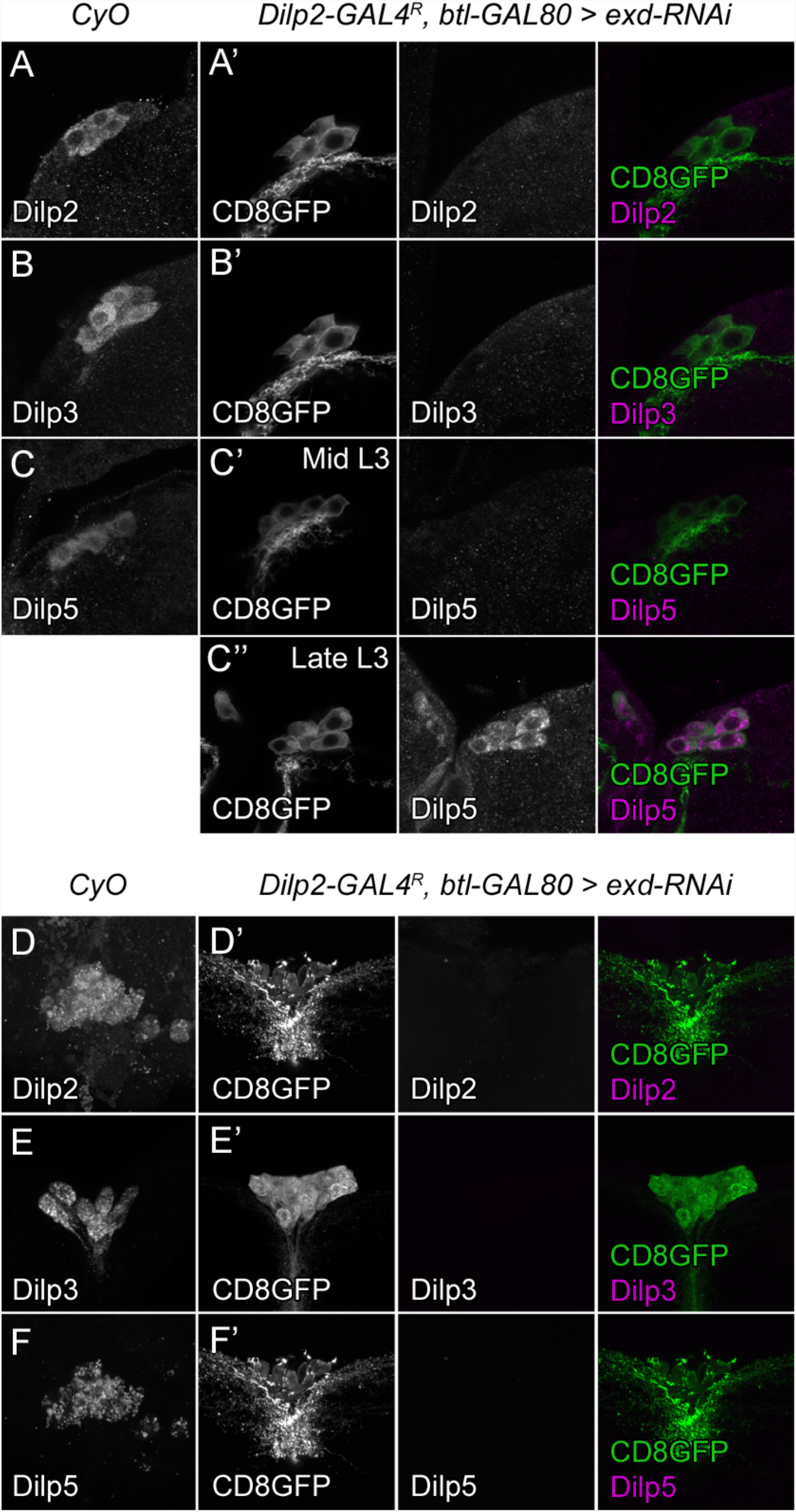
Knockdown of *exd* in the IPCs reduces Dilp protein levels, similar to *hth* knockdown. (A-F’) Genetic depletion of *exd* in the adult or L3 IPCs using Dilp2-GAL4^R^;btl-GAL80 resulted in reduced (A, D) Dilp2, (B, E) Dilp3, and (C, F) Dilp5 protein levels compared to control siblings reared in identical conditions with *exd*^*RNAi*^ #1. Dilp5 was only affected until mid-L3, after which there was no longer any observed effect in late L3 larvae (C”). Flies were reared and age-controlled as described above, then brains from knockdown and balanced sibling flies were dissected and stained in the same tube. Expression of *CD8GFP* was used to differentiate control and test genotypes, and label IPCs. Images were acquired using identical confocal settings.

**Supplemental File 1:** Average dilp2, dilp3, and dilp5 expression values (FPKM) from Huang et al^30^ per DGRP line for males and females, used as the input for the GWAS (http://dgrp2.gnets.ncsu.edu/).

**Supplemental File 2: Adult IPC transcriptome**. Transcriptome analysis of manually sorted adult IPCs. Expression levels are expressed as transcripts per million (TPM).

**Supplemental File 3:** List of all genes detected in the screen that were tested for a role in *dilp* regulation with their corresponding RNAi-line number (BDSC) and adult IPC expression (TPM).

## References

1. Brogiolo, W. et al. An evolutionarily conserved function of the Drosophila insulin receptor and insulin-like peptides in growth control. Curr. Biol. 11, 213–221 (2001).

2. Grönke, S., Clarke, D.-F., Broughton, S., Andrews, T. D. & Partridge, L. Molecular Evolution and Functional Characterization of Drosophila Insulin-Like Peptides. PLOS Genet. 6, e1000857 (2010).

3. Giannakou, M. E. & Partridge, L. Role of insulin-like signalling in Drosophila lifespan. Trends Biochem. Sci. 32, 180–188 (2007).

4. Itskov, P. & Ribeiro, C. The Dilemmas of the Gourmet Fly: The Molecular and Neuronal Mechanisms of Feeding and Nutrient Decision Making in Drosophila. Front. Neurosci. 7, 12 (2013).

5. Vitali, V., Horn, F. & Catania, F. Insulin-like signaling within and beyond metazoans. Biol. Chem. 399, 851–857 (2018).

6. Garofalo, R. S. Genetic analysis of insulin signaling in Drosophila. Trends Endocrinol. Metab. 13, 156–162 (2002).

7. Garelli, A. et al. Dilp8 requires the neuronal relaxin receptor Lgr3 to couple growth to developmental timing. Nat. Commun. 6, 8732 (2015).

8. Vallejo, D. M., Juarez-Carreno, S., Bolivar, J., Morante, J. & Dominguez, M. A brain circuit that synchronizes growth and maturation revealed through Dilp8 binding to Lgr3. Science 350, aac6767 (2015).

9. Imambocus, B. N. et al. A neuropeptidergic circuit gates selective escape behavior of Drosophila larvae. Curr. Biol. 32, 149–163 (2022).

10. Castellanos, M. C., Tang, J. C. Y. & Allan, D. W. Female-biased dimorphism underlies a female-specific role for post-embryonic Ilp7 neurons in Drosophila fertility. Development 140, 3915–3926 (2013).

11. Ohhara, Y., Kobayashi, S., Yamakawa-Kobayashi, K. & Yamanaka, N. Adult-specific insulin-producing neurons in Drosophila melanogaster. J. Comp. Neurol. 526, 1351–1367 (2018).

12. Liu, Y., Liao, S., Veenstra, J. A. & Nassel, D. R. Drosophila insulin-like peptide 1 (DILP1) is transiently expressed during non-feeding stages and reproductive dormancy. Sci. Rep. 6, 26620 (2016).

13. Slaidina, M., Delanoue, R., Gronke, S., Partridge, L. & Leopold, P. A Drosophila insulin-like peptide promotes growth during nonfeeding states. Dev. Cell 17, 874–884 (2009).

14. Post, S. et al. Drosophila insulin-like peptide dilp1 increases lifespan and glucagon-like Akh expression epistatic to dilp2. Aging Cell 18, e12863 (2019).

15. Okamoto, N. & Nishimura, T. Signaling from Glia and Cholinergic Neurons Controls Nutrient-Dependent Production of an Insulin-like Peptide for Drosophila Body Growth. Dev. Cell 35, 295–310 (2015).

16. Rulifson, E. J., Kim, S. K. & Nusse, R. Ablation of insulin-producing neurons in flies: growth and diabetic phenotypes. Science 296, 1118–1120 (2002).

17. Söderberg, J. A. E., Birse, R. T. & Nässel, D. R. Insulin Production and Signaling in Renal Tubules of Drosophila Is under Control of Tachykinin-Related Peptide and Regulates Stress Resistance. PLoS One 6, e19866 (2011).

18. Veenstra, J. A., Agricola, H.-J. & Sellami, A. Regulatory peptides in fruit fly midgut. Cell Tissue Res. 334, 499–516 (2008).

19. Post, S. et al. Drosophila Insulin-Like Peptides DILP2 and DILP5 Differentially Stimulate Cell Signaling and Glycogen Phosphorylase to Regulate Longevity. Front. Endocrinol. (Lausanne). 9, 245 (2018).

20. Kannan, K. & Fridell, Y.-W. C. Functional implications of Drosophila insulin-like peptides in metabolism, aging, and dietary restriction. Front. Physiol. 4, 288 (2013).

21. Nässel, D. R., Kubrak, O. I., Liu, Y., Luo, J. & Lushchak, O. V. Factors that regulate insulin producing cells and their output in Drosophila. Front. Physiol. 4, 252 (2013).

22. Colombani, J. et al. A nutrient sensor mechanism controls Drosophila growth. Cell 114, 739–749 (2003).

23. Agrawal, N. et al. The Drosophila TNF Eiger Is an Adipokine that Acts on Insulin-Producing Cells to Mediate Nutrient Response. Cell Metab. 23, 675–684 (2016).

24. Luo, J., Lushchak, O. V, Goergen, P., Williams, M. J. & Nässel, D. R. Drosophila insulin-producing cells are differentially modulated by serotonin and octopamine receptors and affect social behavior. PLoS One 9, e99732 (2014).

25. Mackay, T. F. C. et al. The Drosophila melanogaster Genetic Reference Panel. Nature 482, 173–178 (2012).

26. Shorter, J. et al. Genetic architecture of natural variation in Drosophila melanogaster aggressive behavior. Proc. Natl. Acad. Sci. U. S. A. 112, E3555–63 (2015).

27. Zwarts, L. et al. The genetic basis of natural variation in mushroom body size in Drosophila melanogaster. Nat. Commun. 6, 10115 (2015).

28. Baker, B. M., Carbone, M. A., Huang, W., Anholt, R. R. H. & Mackay, T. F. C. Genetic basis of variation in cocaine and methamphetamine consumption in outbred populations of Drosophila melanogaster. Proc. Natl. Acad. Sci. U. S. A. 118, e2104131118 (2021).

29. Ivanov, D. K. et al. Longevity GWAS Using the Drosophila Genetic Reference Panel. J. Gerontol. A. Biol. Sci. Med. Sci. 70, 1470–1478 (2015).

30. Huang, W. et al. Genetic basis of transcriptome diversity in Drosophila melanogaster. Proc. Natl. Acad. Sci. U. S. A. 112, E6010–9 (2015).

31. Clements, J., Hens, K., Francis, C., Schellens, A. & Callaerts, P. Conserved role for the Drosophila Pax6 homolog Eyeless in differentiation and function of insulin-producing neurons. Proc. Natl. Acad. Sci. U. S. A. 105, 16183–8 (2008).

32. Okamoto, N., Nishimori, Y. & Nishimura, T. Conserved role for the Dachshund protein with Drosophila Pax6 homolog Eyeless in insulin expression. Proc. Natl. Acad. Sci. 109, 2406–2411 (2012).

33. Kumar, J. P. Retinal determination the beginning of eye development. Curr. Top. Dev. Biol. 93, 1–28 (2010).

34. Buhler, K. et al. Growth control through regulation of insulin signalling by nutrition-activated steroid hormone in Drosophila. Development 145, dev165654 (2018).

35. Abu-Shaar, M., Ryoo, H. D. & Mann, R. S. Control of the nuclear localization of Extradenticle by competing nuclear import and export signals. Genes Dev. 13, 935–945 (1999).

36. Broughton, S. J. et al. Longer lifespan, altered metabolism, and stress resistance in Drosophila from ablation of cells making insulin-like ligands. Proc. Natl. Acad. Sci. U. S. A. 102, 3105–3110 (2005).

37. Zhang, X. et al. Pax6 is regulated by Meis and Pbx homeoproteins during pancreatic development. Dev. Biol. 300, 748–757 (2006).

38. Owa, T. et al. Meis1 Coordinates Cerebellar Granule Cell Development by Regulating Pax6 Transcription, BMP Signaling and Atoh1 Degradation. J. Neurosci. 38, 1277–1294 (2018).

39. Cao, J. et al. Insight into insulin secretion from transcriptome and genetic analysis of insulin-producing cells of Drosophila. Genetics 197, 175–192 (2014).

40. Park, S. et al. A Genetic Strategy to Measure Circulating Drosophila Insulin Reveals Genes Regulating Insulin Production and Secretion. PLoS Genet. 10, e1004555 (2014).

41. Broughton, S. et al. Reduction of DILP2 in Drosophila triages a metabolic phenotype from lifespan revealing redundancy and compensation among DILPs. PLoS One 3, e3721–e3721 (2008).

42. Clements, J., Hens, K., Francis, C., Schellens, A. & Callaerts, P. Conserved role for the Drosophila Pax6 homolog Eyeless in differentiation and function of insulin-producing neurons. Proc. Natl. Acad. Sci. U. S. A. 105, 16183–16188 (2008).

43. Bessa, J., Gebelein, B., Pichaud, F., Casares, F. & Mann, R. S. Combinatorial control of Drosophila eye development by eyeless, homothorax, and teashirt. Genes Dev. 16, 2415–2427 (2002).

44. Lopes, C. S. & Casares, F. hth maintains the pool of eye progenitors and its downregulation by Dpp and Hh couples retinal fate acquisition with cell cycle exit. Dev. Biol. 339, 78–88 (2010).

45. Wang, S., Tulina, N., Carlin, D. L. & Rulifson, E. J. The origin of islet-like cells in Drosophila identifies parallels to the vertebrate endocrine axis. Proc. Natl. Acad. Sci. U. S. A. 104, 19873–19878 (2007).

46. Post, S. & Tatar, M. Nutritional Geometric Profiles of Insulin/IGF Expression in Drosophila melanogaster. PLoS One 11, e0155628 (2016).

47. Peiris, H. et al. Discovering human diabetes-risk gene function with genetics and physiological assays. Nat. Commun. 9, 3855 (2018).

48. Huang, W. et al. Natural variation in genome architecture among 205 Drosophila melanogaster Genetic Reference Panel lines. Genome Res. 24, 1193–1208 (2014).

49. Nagoshi, E. et al. Dissecting differential gene expression within the circadian neuronal circuit of Drosophila. Nat. Neurosci. 13, 60–68 (2010).

50. Chen, S., Zhou, Y., Chen, Y. & Gu, J. fastp: an ultra-fast all-in-one FASTQ preprocessor. Bioinformatics 34, i884–i890 (2018).

51. Patro, R., Duggal, G., Love, M. I., Irizarry, R. A. & Kingsford, C. Salmon provides fast and bias-aware quantification of transcript expression. Nat. Methods 14, 417–419 (2017).

52. Soneson, C., Love, M. I. & Robinson, M. D. Differential analyses for RNA-seq: transcript-level estimates improve gene-level inferences. F1000Research 4, 1521 (2015).

53. Schindelin, J. et al. Fiji: an open-source platform for biological-image analysis. Nat. Methods 9, 676–682 (2012).

54. Schneider, C. A., Rasband, W. S. & Eliceiri, K. W. NIH Image to ImageJ: 25 years of image analysis. Nat. Methods 9, 671–675 (2012).

55. Bürgy-Roukala, E., Miellet, S., Mishra, A. K. & Sprecher, S. G. Early Eye Development: Specification and Determination BT - Molecular Genetics of Axial Patterning, Growth and Disease in the Drosophila Eye. in (eds. Singh, A. & Kango-Singh, M.) 1–36 (Springer New York, 2013). doi:10.1007/978-1-4614-8232-1_1.

